# Novel insights on the contribution of plastoglobules and reactive oxygen species to chromoplast differentiation

**DOI:** 10.1101/2022.06.20.496796

**Authors:** Luca Morelli, Salvador Torres-Montilla, Gaetan Glauser, Venkatasalam Shanmugabalaji, Felix Kessler, Manuel Rodriguez-Concepcion

## Abstract

Enriching plant tissues in phytonutrients can be done by stimulating their biosynthesis but also by providing appropriate sink structures for their sequestering and storage. Chromoplasts are plastids specialized in the production and accumulation of carotenoids that are naturally formed in non-photosynthetic tissues such as flower petals and ripe fruit. Chromoplasts can also be artificially differentiated from leaf chloroplasts by boosting the production of phytoene (the first committed intermediate of the carotenoid pathway) with the bacterial phytoene synthase crtB. Here we show that crtB-induced leaf chromoplasts develop plastoglobules harboring high levels of carotenoids (mainly phytoene and pro-vitamin A β-carotene) but also other nutritionally-relevant isoprenoids such as tocopherols (vitamin E) and phylloquinone (vitamin K1). Further promoting plastoglobule proliferation by exposure to intense (high) light resulted in a higher accumulation of these health-related metabolites but also an acceleration of the chloroplast-to-chromoplast conversion. We further show that production of reactive oxygen species (ROS) stimulates chromoplastogenesis. Our data suggest that, similar to that already described for decreased photosynthesis and enhanced carotenoid biosynthesis, ROS production is not just a consequence but a promoter of the chromoplast differentiation process.

## Introduction

Without regular and nutritious food, humans cannot live, fend off diseases or lead productive lives. Malnutrition actually goes beyond low food intake, as more than two billion adults, adolescents and children are now obese or overweight according to the Food and Agriculture Organization, FAO (http://www.fao.org/sustainable-development-goals/goals/goal-2/en/). The consequences are severe for public health and for individuals’ and communities’ quality of life. A powerful tool to prevent micronutrient deficiency and achieve optimal nutrition is biofortification, i.e., improving the production and accumulation of health-promoting bioactive metabolites in plants and other food sources. While biofortification offers advantages over food fortification (i.e., adding micronutrients to harvested or processed food) or dietary supplements, breeding and biotechnological approaches aimed to boost the production and accumulation of phytonutrients in crops are still underdeveloped, partially due to our limited knowledge of how plants regulate the biosynthesis and storage of these metabolites.

Plant isoprenoids include a wide diversity of metabolites that humans cannot produce but must acquire from food sources. In particular, many plastidial isoprenoids with roles in photosynthesis and photoprotection also serve in animal cells as vitamins and powerful antioxidants (Muñoz and Munné-Bosch, 2019; Rodriguez-Concepcion et al., 2018; Fitzpatrick et al., 2012). They include carotenoids such as β-carotene (the main precursor of vitamin A), chromanols such as tocopherols (vitamin E) and plastochromanol, and quinones such as phylloquinone (vitamin K1), plastoquinone and the tocopherol oxidation product α-tocopherolquinone or α-tocoquinone (Figure 1). These nutritionally relevant plastidial isoprenoids derive from precursors produced by the methylerythritol 4-phosphate (MEP) pathway. Chromanols and quinones also include moieties derived from the shikimate pathway (Figure 1). All three classes of isoprenoid metabolites are present in different plastid types, where they have different roles. In chloroplasts they are mainly located in photochemically active thylakoid membranes (where they participate in photosynthesis-related processes) and, to a lower extent, in the envelope (Lichtenthaler, 2012). Notably, tocopherols are particularly abundant in the gerontoplasts that differentiate from chloroplasts in senescent leaves, whereas carotenoids accumulate at highest levels in the chromoplasts present in non-photosynthetic pigmented tissues such as flower petals and ripe fruits. Excess amounts of carotenoids, chromanols and quinones accumulate in thylakoid-derived lipid bodies or lipoprotein particles known as plastoglobules (PG), which are normally produced under a wide range of stress and developmental conditions causing a disassembly of the thylakoid membrane system (Van Wijk and Kessler, 2017). PG proliferate in both gerontoplasts and chromoplasts (Berry et al., 2019; Fraser et al., 2007; Nogueira et al., 2013; Muñoz and Munné-Bosch, 2019; Sadali et al., 2019; Sun et al., 2018; Van Wijk and Kessler, 2017). Besides acting as storage compartments, PG also appear to play a role in the production of chromanols and quinones in chloroplasts (Van Wijk and Kessler, 2017). The presence of several carotenoid biosynthetic enzymes in the PG proteome of pepper ripe fruit (Ytterberg et al., 2006) (Figure 1) also suggests that PG are more than just storage sites for carotenoids in chromoplasts (Van Wijk and Kessler, 2017).

**Figure 1.**
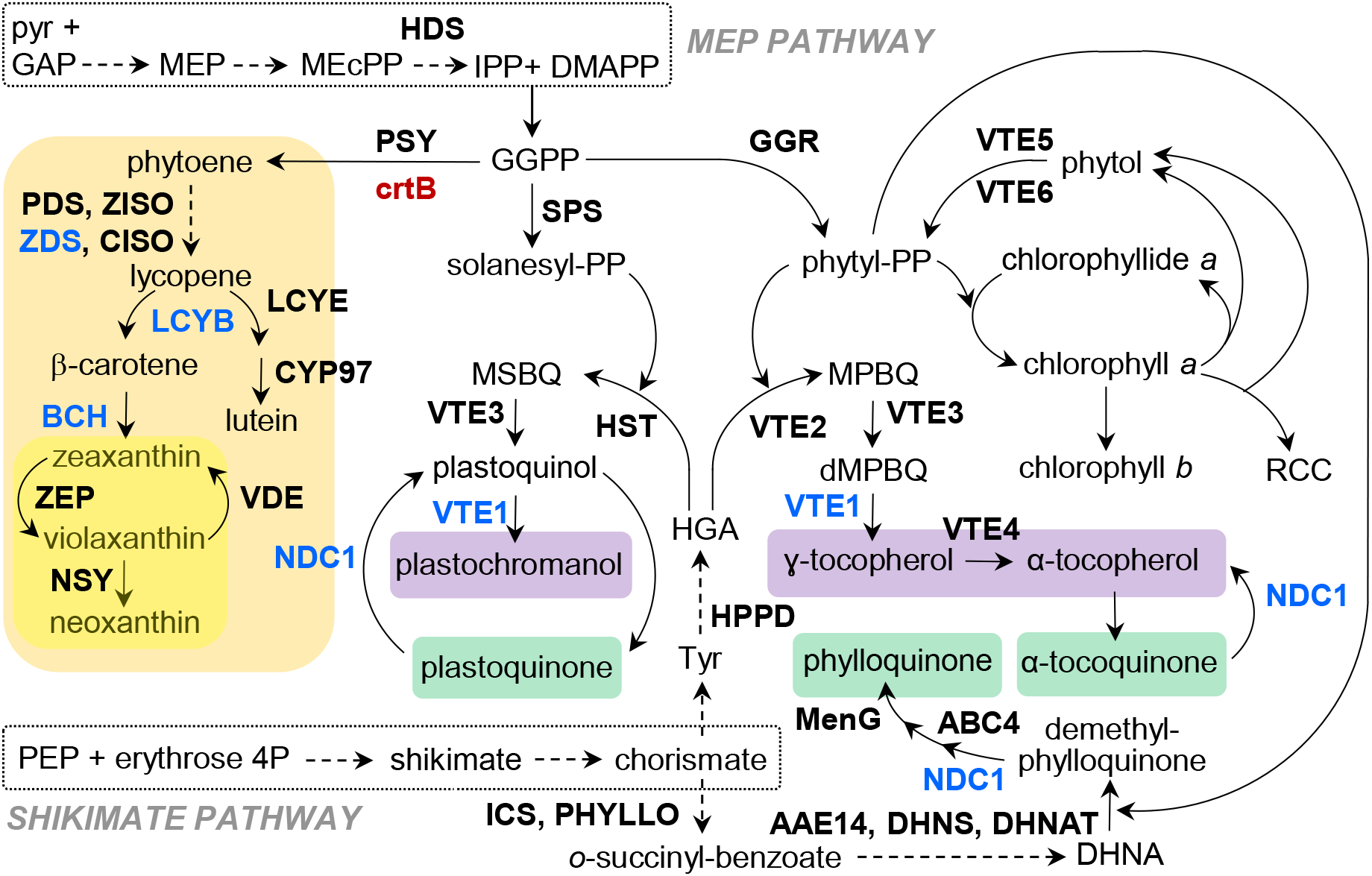
Schematic plastidial isoprenoid pathways. Carotenoids are boxed in orange (with β,β-xanthophylls marked in yellow), chromanols in purple, and quinones in green. Dashed arrows represent multiple steps. Enzymes are shown in bold type, and those found in plastoglobules (PG) are indicated in blue. Accession numbers are listed in Table S1.

Extensive genetic evidence indicates that an increased sink capacity leads to enhanced levels of plastidial isoprenoids such as carotenoids and tocopherols, but our limited knowledge of how plastid differentiation is regulated has prevented to fully exploit this “pull” strategy as an alternative or a complement to the “push” approach based on stimulating biosynthesis (Garcia-Mas and Rodriguez-Concepcion, 2016). In particular, surprisingly little is known on the factors involved in natural differentiation of chloroplasts into chromoplasts (Sadali et al., 2019; Torres-Montilla and Rodriguez-Concepcion, 2021). We have recently demonstrated that chromoplasts harboring high levels of endogenous carotenoids can be synthetically created in leaves and other chloroplast-harboring organs of any plant species by overexpressing the *Pantoea ananatis crtB* gene, encoding a bacterial phytoene synthase (Houhou et al., 2022; Llorente et al., 2020). Plastid targeting of either the unmodified crtB protein (by means of a cryptic plastid targeting sequence present in the protein) or of a modified version with a plant plastid targeting sequence fused to its N-terminus (referred to as (p)crtB) resulted in a dramatic production of phytoene, the first committed intermediate of the carotenoid pathway (Figure 1). This phytoene boost was proposed to interfere with photosynthesis and hence precondition chloroplasts to become chromoplasts as phytoene was later converted into downstream carotenoids by the endogenous plant enzymes (Llorente et al., 2020). This work confirmed that down-regulation of photosynthesis and up-regulation of carotenoid accumulation were not just consequences but requirements for chromoplastogenesis. It further provided a new model system to study the differentiation of chromoplasts without the interference of developmental or environmental cues that normally regulate this process in nature (e.g., during fruit ripening).

Agroinfiltration of tobacco (*Nicotiana benthamiana*) leaves with crtB or (p)crtB constructs results in a massive reprogramming of nuclear gene expression and global cell metabolism to support the profound changes that occur in plastid ultrastructure as endogenous chloroplasts transform into artificial chromoplasts. In particular, crtB-induced leaf chromoplasts developed tightly appressed membrane stacks and showed a proliferation of round structures tentatively identified as PG (Llorente et al., 2020). Using the (p)crtB version, from herein referred to as crtB, here we provide a deep characterization of these subplastidial structures and show that they accumulate high levels of carotenoids but also other nutritionally-relevant isoprenoids such as chromanols and quinones. We also demonstrate that the PG that proliferate in artificial leaf chromoplasts host active crtB enzymes and accumulate virtually all crtB-derived phytoene as well as most of the β-carotene produced in the tissue, supporting a role for these lipid bodies in carotenoid biosynthesis. Further promoting PG development by pre-exposing leaves to intense (high) light resulted in a higher accumulation of carotenoids, tocopherols and phylloquinone but also an acceleration of the chloroplast-to-chromoplast differentiation process. Finally, we show that redox changes that take place as chloroplasts transform into chromoplasts positively contribute to the differentiation process.

## Results and Discussion

### Phytoene production by crtB triggers disorganization of photosynthetic complexes followed by proliferation of PG as leaf chloroplasts turn into chromoplasts

Transmission electron microscopy (TEM) images of crtB-induced leaf chromoplasts show characteristic features such as electron-dense membrane stacks and PG-resembling structures (Figure 2). By contrast, chloroplasts present in control leaves expressing GFP are characterized by a regular network of thylakoidal membranes and grana as well as a low number of PG (Llorente et al., 2020). To validate these observations with quantitative data, TEM images from tobacco leaves agroinfiltrated with either GFP or crtB constructs were used for image analysis. First, we measured grana lateral irregularity (GLI), defined as the coefficient of variation (the ratio of the standard deviation to the mean) of the length of each membrane layer forming the granum (in chloroplasts) or the membrane stack (in chromoplasts) (Kowalewska et al., 2016). In an ideal situation of chloroplast grana, which have a cylindrical shape, this value is around 0. By contrast, in a situation of complete irregularity the value is near 1. In our analysis, the GLI value for GFP samples was lower than 0.1 (indicating a certain regularity in grana typical of chloroplasts) whereas a much higher GLI value was observed for crtB samples, illustrating the disorganization of internal membranes observed in the differentiated chromoplasts (Figure 2A).

**Figure 2.**
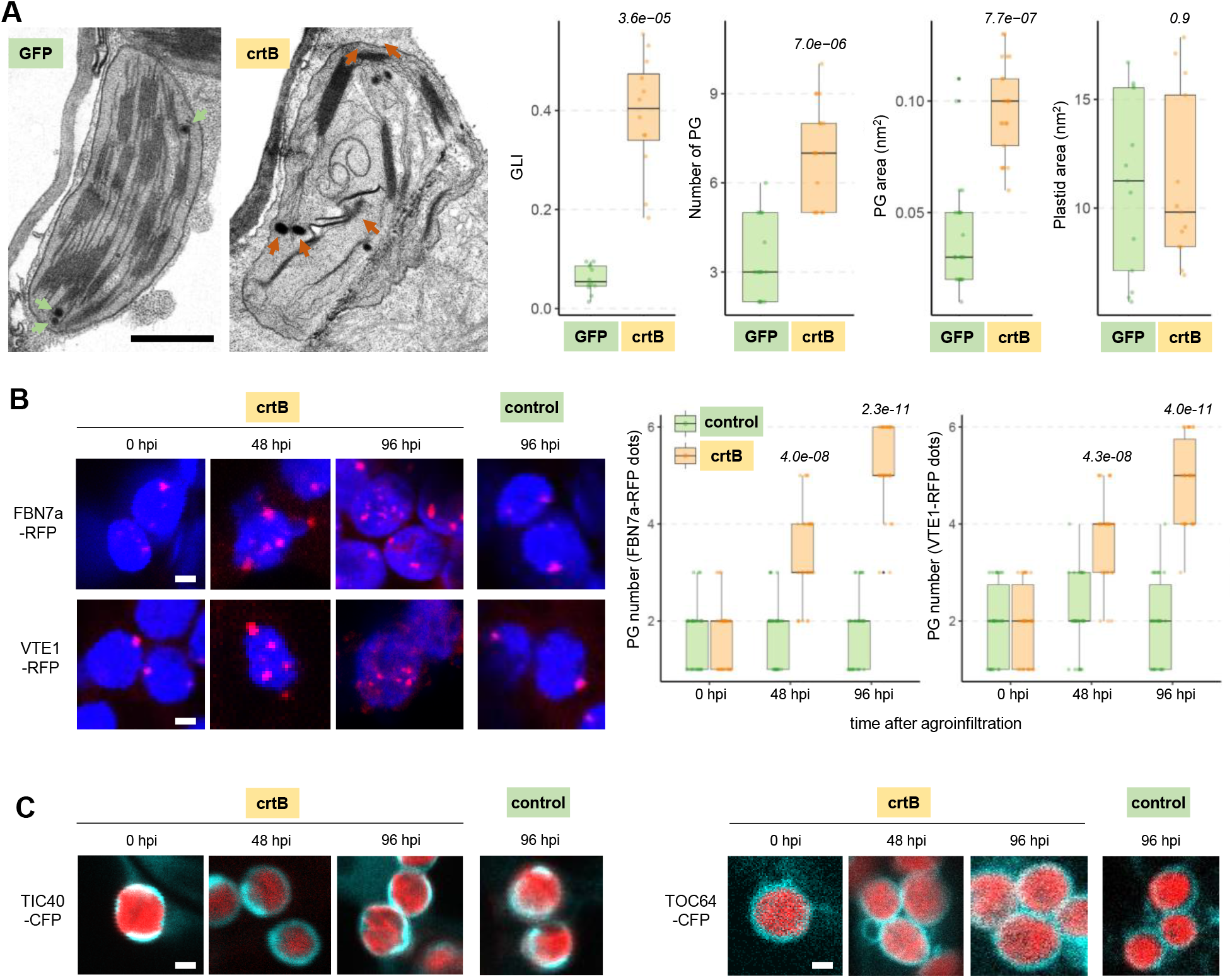
Agroinfiltration with crtB generates artificial chromoplasts with a proliferation of PG. (A) TEM images of representative plastids from *N. benthamiana* leaves agroinfiltrated with constructs harboring crtB or a GFP control, as indicated, and collected at 120 hpi. PG are marked with arrows. Boxplots represent quantitative data on grana lateral irregularity (GLI), PG number, PG area and total plastid area collected from TEM images of n=12 chloroplasts (GFP) or chromoplasts (crtB) from different plants. Boxes represent the values between the upper and the lower quartile, horizontal line the median, whiskers the upper and lower values, and dots the individual values (the ones outside the whiskers represent outliers). Numbers in italics indicate Wilkinson test *p* values. (B) Localization of the indicated PG markers in plastids at the indicated time points after co-agroinfiltration with crtB or not (control). RFP fluorescence from PG markers is shown in red, and chlorophyll autofluorescence in blue. Boxplots are like those in (A) and represent quantitative data collected from confocal images of n=30 plastids for each set of constructs. Numbers in italics indicate Kruskal-Wallis test *p* values. (C) Localization of markers of the inner or outer envelope membranes (TIC40-CFP or TOC64-CFP, respectively) in plastids at the indicated time points after co-agroinfiltration with crtB or not (control). CFP fluorescence is shown in turquoise, and chlorophyll autofluorescence in red. All pictures of the same section are to the same magnification. Bar size, 1 µm.

We next measured the number and area of electron-dense bodies (PG) in the plastids from GFP and crtB samples and confirmed that the artificial chromoplasts that developed upon expression of crtB contained PG that were larger and more abundant than those in the chloroplasts of GFP controls (Figure 2A). These results were further validated using PG marker proteins fused to RFP and expressed under the control of glucocorticoid-induced promoters (Figure 2B). The rationale of using inducible promoters was to prevent protein overproduction and aggregation but also putative interferences with the differentiation process arising from the constitutive presence of high levels of PG-associated proteins such as the fibrillin FBN7a and the chromanol biosynthetic enzyme VTE1 (Figure 1) (Van Wijk and Kessler, 2017). Tobacco leaves were agroinfiltrated with the PG marker protein constructs either alone or together with untagged crtB. Localization of the fluorescent marker proteins was analyzed by confocal laser scanning microscopy (CLSM) 24 h after inducing their expression with β-estradiol. In chloroplasts of control leaves, only a few fluorescent dots corresponding to PG were observed, and their number did not change up to 96 hours post-infiltration (hpi). When crtB was co-expressed with the marker proteins, however, the number of plastidial dots increased with time (i.e., as chromoplasts developed from chloroplasts), more than doubling their numbers compared to control samples at 96 hpi (Figure 2B). In similar experiments using inducible CFP-fused marker proteins of the envelope outer and inner membranes (TOC64 and TIC40, respectively), the localization of the markers was similar in control and crtB samples at all stages (Figure 2C). These results, together with the absence of vesicle-like structures forming from inner envelope membranes in TEM images (Figure 2A) (Llorente et al., 2020), argue against the envelope membranes as major contributors to the newly formed internal membrane systems that develop in crtB-triggered chromoplasts. Instead, it is most likely that the disintegration of thylakoid membranes is responsible for the formation of the dense membrane stacks and proliferating PG observed in these plastids.

To investigate the dynamics of the observed membrane reorganization, we followed the levels of marker proteins of plastidial subcompartments by immunoblot analysis of tobacco leaves at several timepoints after agroinfiltration (Figure 3). For these experiments we triggered chromoplastogenesis using crtB-GFP instead of crtB for visualization of the fusion protein by immunoblot analysis using anti-GFP antibodies. The crtB-GFP protein was hardly detectable up to 36 hpi, but later its levels increased steadily, a profile that was expectedly similar to that of its product, phytoene (Figure 3A). Effective quantum yield of PSII (ɸPSII) dropped as phytoene increased, further confirming that the crtB-GFP fusion is an active enzyme able to produce enough phytoene to challenge photosynthesis in leaf chloroplasts and induce their conversion into chromoplasts (Figure 3A). The levels of some but not all proteins associated to photosynthetic membranes were decreased as chromoplasts developed (Figure 3A-B). LHCB2, a component of the light-harvesting complexes, did not change in the 96 hpi period investigated. By contrast, the levels of PsaD (a component of PSI) and PsbA (a component of PSII) dropped between 36 and 48 hpi. An even earlier decrease was detected in the case of the cytochrome b6-f complex subunit PetC, which unlike PsbA and PsaD became undetectable at 96 hpi (Figure 3A-B). The differences in protein levels observed in crtB-GFP samples at 96 hpi compared to 0 hpi (Figure 3B) were very similar to those detected when compared to GFP controls at 96 hpi (Figure 3C), confirming that they illustrate chromoplast vs. chloroplast differences. Strikingly, comparison of transcript levels in GFP and crtB samples at 96 hpi (Llorente et al., 2020) showed very different trends of gene expression and protein accumulation (Figure 3D and Figure S1). The expression of genes encoding LHCB2 was significantly reduced in artificial chromoplast despite protein levels did not change, whereas those for PsaD, PsbA and PetC did not show significant changes that could explain the observed changes in protein accumulation with perhaps the only exception of PsaD (Figure 3). These data together indicate that the early burst in phytoene production that takes in *crtB*-expressing leaves is linked with a drop in ɸPSII likely derived from the transcription-independent down-regulation of proteins associated with photosynthetic electron transport that are encoded either by the nuclear genome (PsaD and PetC) or by the plastome (PsbA).

**Figure 3.**
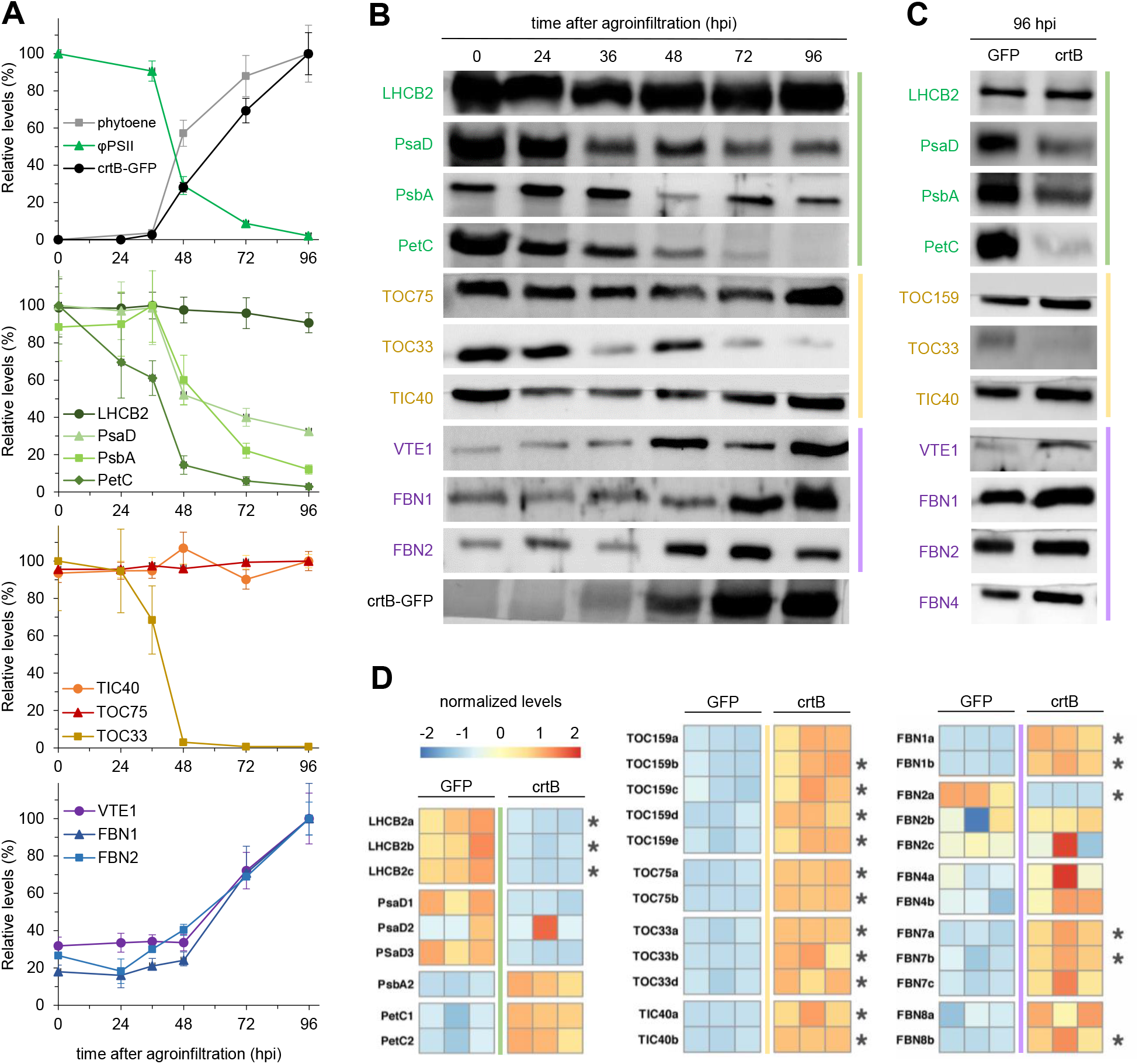
Leaf chromoplast differentiation involves changes in the plastid proteome. Colors represent different subplastidial localizations: thylakoids (green), envelope (orange/red) and PG (purple/blue). (A) Quantification of phytoene, φPSII and protein levels at the indicated points after agroinfiltration with a plastid-targeted GFP-tagged crtB contruct (crtB-GFP). Data are represented relative to the levels just before agroinfiltration (0 hpi) and correspond to the mean and standard deviation of n=5 different leaf samples. (B) Representative immunoblot analysis used to calculate proteins level values shown in (A). (D) Immunoblot analysis of the indicated proteins in leaf samples at 96h after agroinfiltration with either GFP or crtB constructs. (C) Heatmap representing transcript abundance of the indicated genes at 96 hpi in GFP and crtB samples. Data were retrieved from previously reported RNA-seq analysis (Llorente et al., 2020). Columns correspond to replicates. Levels are normalized according to the row_scale of the pheatmap package. Asterisks mark differentially expressed genes when comparing mean crtB vs. GFP values (DESeq2, FDR ≤ 0.05). Gene accessions are available in Table S2.

The plastome-encoded PSII reaction center protein PsbA, also known as D1, is prone to photodamage and further degradation, and it must be replaced by nascent D1 to maintain photosynthetic activity (Järvi et al., 2015). Phytoene build-up might interfere with photoprotection (hence causing a higher rate of D1 degradation) or/and with the production of new D1 proteins, eventually disrupting the PSII repair cycle. In the case of nuclear-encoded PsaD and PetC, their reduced levels could be due to decreased import or increased degradation within the plastid. Import of most plastidial proteins takes place through translocon complexes located at the outer (TOC) and inner (TIC) envelope membranes. While the TIC protein TIC40 showed similar levels from 0 to 96 hpi, the TOC protein TOC33 dropped at 36 hpi (Figure 3B). TOC33 together with TOC159 and TOC75 are major TOC components preferentially involved in the import of photosynthesis-related proteins, and all three of them are targeted for ubiquitination and degradation by the RING-type ubiquitin E3 ligase SP1 in *Arabidopsis thaliana* (Ling et al., 2012). SP1 activity hence modifies the balance between different translocons and the operation of substrate-specific protein import pathways to facilitate plastid differentiation (e.g., the transition of etioplasts to chloroplasts or chloroplasts to gerontoplasts). Recent evidence also indicates that the tomato (*Solanum lycopersicum*) SP1 homologues regulate the conversion of chloroplasts into chromoplasts during fruit ripening (Ling et al., 2021). Up-regulated levels of SP1 proteins in tomato fruit were found to reduce levels of TOC75 but not TIC40, eventually promoting chloroplast to chromoplast transition. However, crtB-dependent chromoplast differentiation in tobacco leaves did not involve changes in the levels of SP1 targets such as TOC75 (Figure 3B) or TOC159 (Figure 3C). Furthermore, expression of *crtB* in Arabidopsis mutant plants defective in SP1 using the viral vector TuMV-crtB (Llorente et al, 2020) resulted in leaves showing the same characteristic chromoplast-associated yellow phenotype and carotenoid overaccumulation observed in wild-type (WT) plants (Figure 4), suggesting that SP1 activity is not required for crtB-mediated chromoplastogenesis.

**Figure 4.**
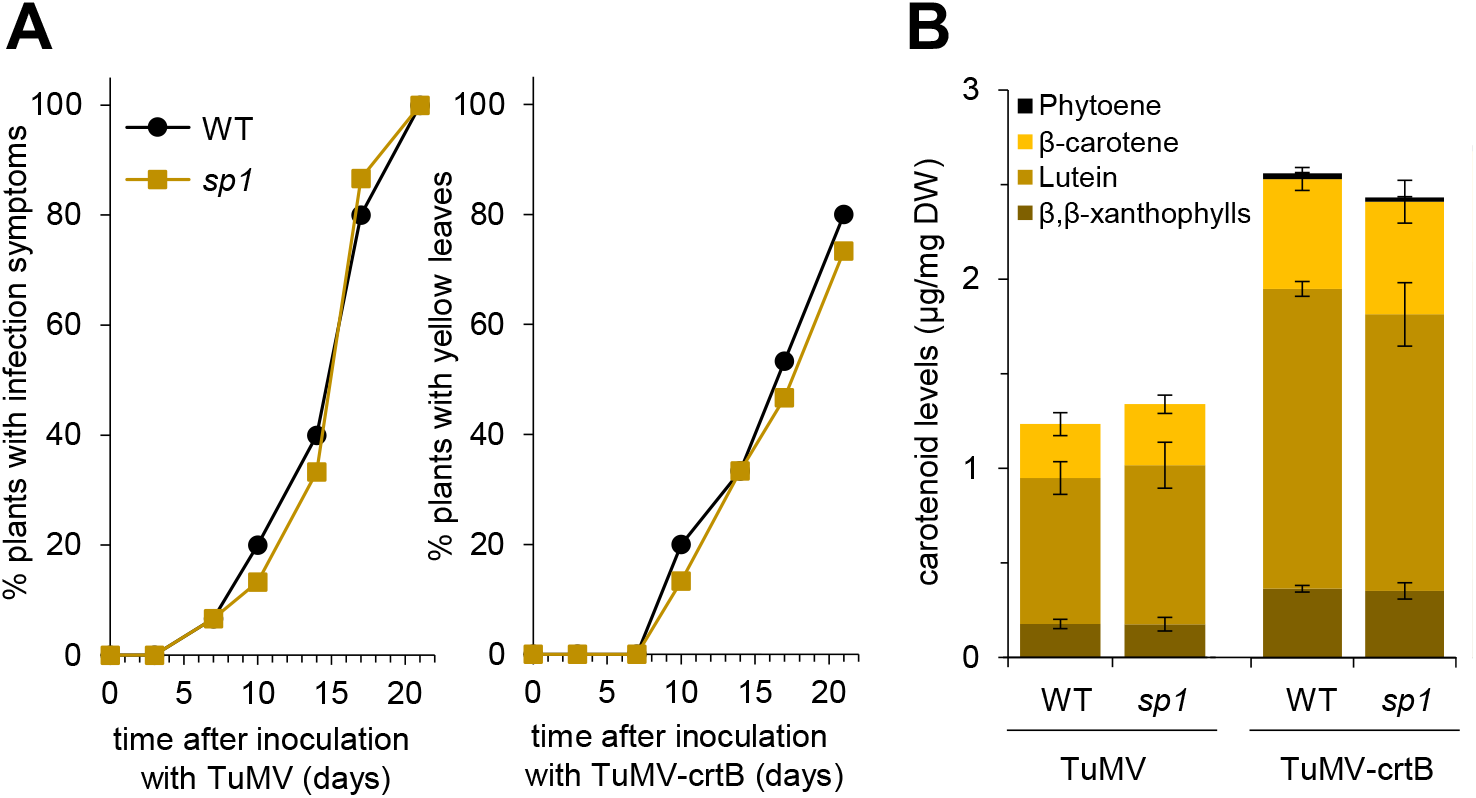
Leaf chromoplastogenesis is not affected in Arabidopsis *sp1* mutants. Soil-grown Arabidopsis plants of the indicated genotypes were infected with viral vector TuMV-crtB or an empty TuMV control and the visual and metabolic phenotypes were monitored afterwards. (A) Rate of development of phenotypes associated with TuMV infection (e.g., curled leaves) or with ctrB activity (e.g., yellow leaves). Data correspond to a representative experiment with n=15 plants. (B) HPLC analysis of carotenoid levels 3 weeks after infection with either TuMV or TuMV-crtB. Data correspond to the mean and standard deviation of n=3 pools of 5 plants each.

At the gene expression level, artificial chromoplastogenesis in *N. benthamiana* leaves resulted in significantly enhanced levels of transcripts for TIC and TOC components, including TOC33 (Figure 3D and Figure S1). These data suggest that post-transcriptional mechanisms tightly regulate the abundance of these important proteins. We speculate that the decrease in TOC33 observed when crtB is expressed in tobacco leaves might be a SP1-independent process that selectively targets this TOC component to prevent import of photosynthetic proteins. In agreement with the conclusion that only the import of photosynthetic proteins might be compromised by reduced TOC33 levels, the levels of nuclear-encoded PG-associated proteins such as VTE1 and fibrillins such as FBN1 or FBN2 did not decrease but increased after 48 hpi (Figure 3B). PG-associated FBN4 also became more abundant in crtB-triggered chromoplasts (Figure 3C). Fibrillins are a family of proteins including PG-associated members (FBN1, FBN2, FBN4, FBN7 and FBN8) that constitute the main portion of the PG proteome and have been attributed mainly structural roles (Van Wijk and Kessler, 2017; Shanmugabalaji et al., 2022, 2013). The expression of *N. benthamiana* genes encoding homologues of PG-associated fibrillins increased in crtB samples at 96 hpi (Figure 3D and Figure S1), suggesting that the observed accumulation of PG-associated proteins was supported, at least in part, by up-regulated gene expression. Together, these data suggest that increased phytoene production by crtB rapidly challenges photosynthesis via down-regulation of the levels of proteins associated with photosynthetic electron transport through mechanisms that might include degradation of PsbA and reduced import of PetC and PsbA (via TOC33). A bit later, an increase in the amounts of PG-associated fibrillins takes place supported by up-regulated gene expression. As a result, PG proliferate likely at the expense of photosynthetic (thylakoid) membranes. Other thylakoids and grana might be packed together to form the membrane stacks observed in leaf chromoplasts, which likely retain chlorophylls (Llorente et al, 2020) and chlorophyll-associated proteins such as LHCB2 (Figure 3) but lack photosynthetic functionality (i.e., the capacity to carry on the electron transport chain).

### Chromoplasts obtained by overexpressing crtB are enriched in lipophilic plastidial isoprenoids besides carotenoids

It is widely known that chromoplasts can store a wide array of lipophilic compounds besides carotenoids, including isoprenoids such as chromanols and quinones. The electron-dense nature of the membrane stacks and PG that developed in crtB-triggered leaf chromoplasts (Figure 2) illustrate their osmiophilicity, i.e., their reactivity with the OsO_4_ present in the TEM sample preparation medium. As this feature is normally associated to unsaturated lipids (Van Wijk and Kessler, 2017), it was concluded that the newly formed structures might sequester enhanced levels of isoprenoid lipids. To test this possibility, we used ultra-high pressure liquid chromatography coupled to mass spectrometry (UHPLC-QTOF-MS) to analyze the levels of isoprenoid chromanols and quinones in tobacco leaves agroinfiltrated with either GFP or crtB and collected at 96 hpi (Figure 5). The results showed that *crtB*-overexpressing leaves were enriched not only in carotenoids but also in total tocopherols (2.5-fold times more than GFP controls), α-tocoquinone (2-fold), phylloquinone (1.7-fold), and plastochromanol-8 (1.8-fold) (Figure 5A). By contrast, the levels of chlorophylls remained unchanged, as previously reported (Llorente et al, 2020), whereas the amount of plastoquinone-9 decreased in artificial chromoplasts (Figure 5A). Ubiquinone, a mitochondrial isoprenoid quinone, showed no changes (Figure 5A), supporting the conclusion that the variations observed in the amount of plastidial isoprenoids are caused by changes occurring at the plastid level. The expression of genes encoding carotenoid and phylloquinone biosynthetic enzymes showed a coordinated increase in crtB compared to GFP leaf areas (Figure 5B and Figure S1). By contrast, no correlation was found between biosynthetic gene expression profiles and metabolite levels in the case of tocopherols or plastoquinone (Figure 5B and Figure S1). We therefore conclude that the improved accumulation of health-related isoprenoids in artificial chromoplasts is not fully supported by biosynthetic gene expression. Instead, the new PG and membrane systems that develop in artificial chromoplasts might substantially improve the catalytic activity of biosynthetic enzymes (e.g., many of which are membrane-associated) or the sequestration capacity of the plastids while protecting the lipophilic isoprenoid metabolites from degradation. To next investigate whether the accumulated isoprenoid metabolites had a differential distribution inside the plastid, we performed a subplastidial membrane fractionation experiment by flotation density centrifugation (Vidi et al., 2006; Perello et al., 2016) (Figure 6). We isolated whole plastids from tobacco leaf tissue at different time points after agroinfiltration with crtB-GFP and then hypertonically lysed them to separate the insoluble (i.e., membrane) fraction, which was resuspended in 45% sucrose. Then, this membrane-containing solution was overlaid with a discontinuous sucrose gradient made with a series of 38%, 20%, 15%, and 5% sucrose (Figure 6A). After ultracentrifugation, fractions were collected and used for immunoblot analyses of protein markers from different subplastidial membrane compartments (Figure 6B). The immunoblot analysis of 0 hpi samples (i.e., immediately before the agro-infiltration, only containing chloroplasts) with an antibody against the Arabidopsis PG-associated FBN1a protein only showed immunoreactive bands in the upper fractions, corresponding to the 5% section. The envelope marker TOC75 was detected in the fractions isolated from lower 38% section but also in those from the 45% section. The thylakoid membrane marker LHCB2 was only detected in the bottom fractions of the 45% section, progressively fading towards the limit with the 38% section (Figure 6B). Based on these and previous results (Vidi et al., 2006; Perello et al., 2016) the next experiments were done by pooling fractions in three groups (Figure 6A). The first group corresponded to the 5% and 15% sections, and it was named PG as it included the fractions where PG are normally located. The second one, referred to as M1, corresponded to the 20% and 38% sections and it contained envelopes and most likely other non-photosynthetic membrane types. The third one, named M2, corresponded to the 45% section and it was enriched in thylakoid and envelope membranes. Samples belonging to these three groups of fractions were collected at different timepoints after agroinfiltration with crtB-GFP and pooled together in a single tube (Figure 6A). Immunoblot analyses of these pools together with stroma fractions and whole plastid extracts confirmed their membrane composition (Figure 6C). We next analyzed the isoprenoid contents of PG, M1 and M2 pools in crtB-GFP samples (Figure 6D). In PG, all isoprenoids measured increased at 48 hpi and then either kept increasing (phytoene, β-carotene, α-tocoquinone, phylloquinone, plastochromanol, α-tocopherol) or decreased (lutein, plastoquinone) at 96 hpi. The increase was most dramatic for carotenoids (44-fold increase at 96 hpi relative to 0 hpi). The PG pool lacked any color at 0 hpi (when carotenoids were almost missing) but it acquired an orange-brownish color and oily consistency at 48 hpi and, most strongly, 96 hpi (Figure 6A), as the levels of the orange carotenoid β-carotene increased (Figure 6D). In the case of M1 membranes, phylloquinone was almost undetectable, plastoquinone and tocopherols were reduced, and only carotenoid contents were raised (23-fold increase at 96 hpi) due to a preferential accumulation of lutein (yellow) and, to a much lower extent, β-carotene. In agreement with this carotenoid profile, the M1 pool was transparent at 0 hpi and it progressively turned yellow as chloroplasts became chromoplasts (Figure 6A). The thylakoid-rich M2 membranes lost their plastoquinone content and showed slightly enhanced levels of tocopherols and carotenoids as chromoplasts differentiated (Figure 6D). However, color changes were not detected because of the presence of chlorophylls (Figure 6A). Together, the newly formed membrane structures in artificial leaf chloroplasts appear to promote the accumulation of plastidial isoprenoids. The case of PG is particularly interesting as their proliferation in these leaf chromoplasts leads to enhanced levels of isoprenoid vitamins A (β-carotene), E (tocopherols and particularly α-tocopherol) and K1 (phylloquinone). The existence of common storage structures for different types of plastidial isoprenoids might explain, at least in part, why engineering the carotenoid pathway (Fraser et al., 2007) or tocopherol biosynthesis (Lu et al., 2013) can result in higher levels of both carotenoids and tocopherols.

**Figure 5.**
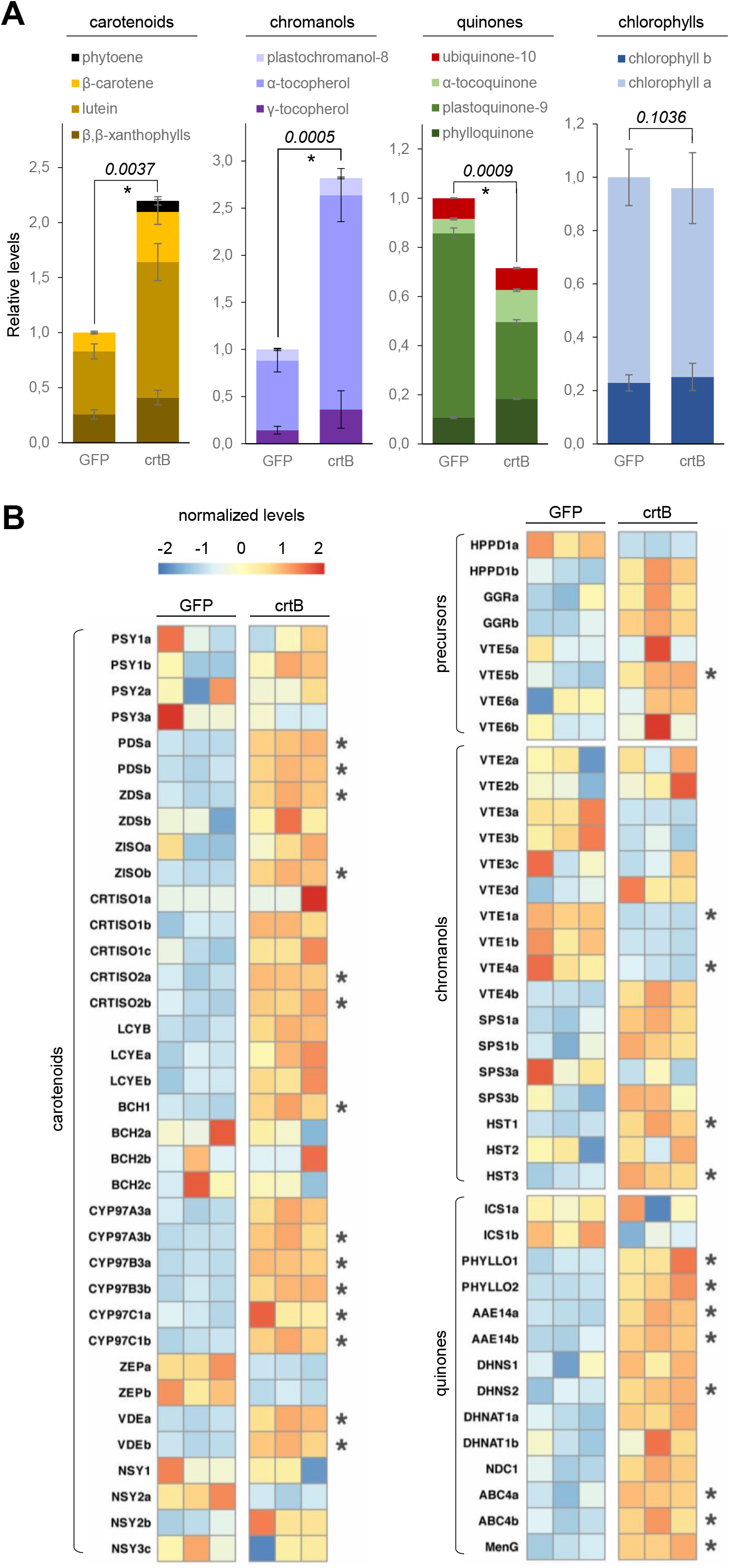
Artificial leaf chromoplasts accumulate high levels of plastidial isoprenoids. (A) Levels of the indicated isoprenoids in *N. benthamiana* leaves agroinfiltrated with constructs harboring crtB or a GFP control at 96 hpi. Data are represented relative to the levels in GFP samples and correspond to the mean and standard deviation of n=3 different leaves. Numbers in italics indicate *p* values of Student *t*-test analyses comparing total levels, and asterisk marks statistically significant differences (*p* ≤ 0.01). (B) Heatmap representing transcript abundance of the indicated isoprenoid biosynthetic genes at 96 hpi. Data were retrieved from previously reported RNA-seq analysis (Llorente et al., 2020). Columns correspond to replicates. Levels are normalized according to the row_scale of the pheatmap package. Asterisks mark differentially expressed genes when comparing mean crtB vs. GFP values (DESeq2, FDR ≤ 0.05). See Figure 1 for the position of the corresponding enzymes in isoprenoid biosynthetic pathways. Gene accessions are available in Table S1.

**Figure 6.**
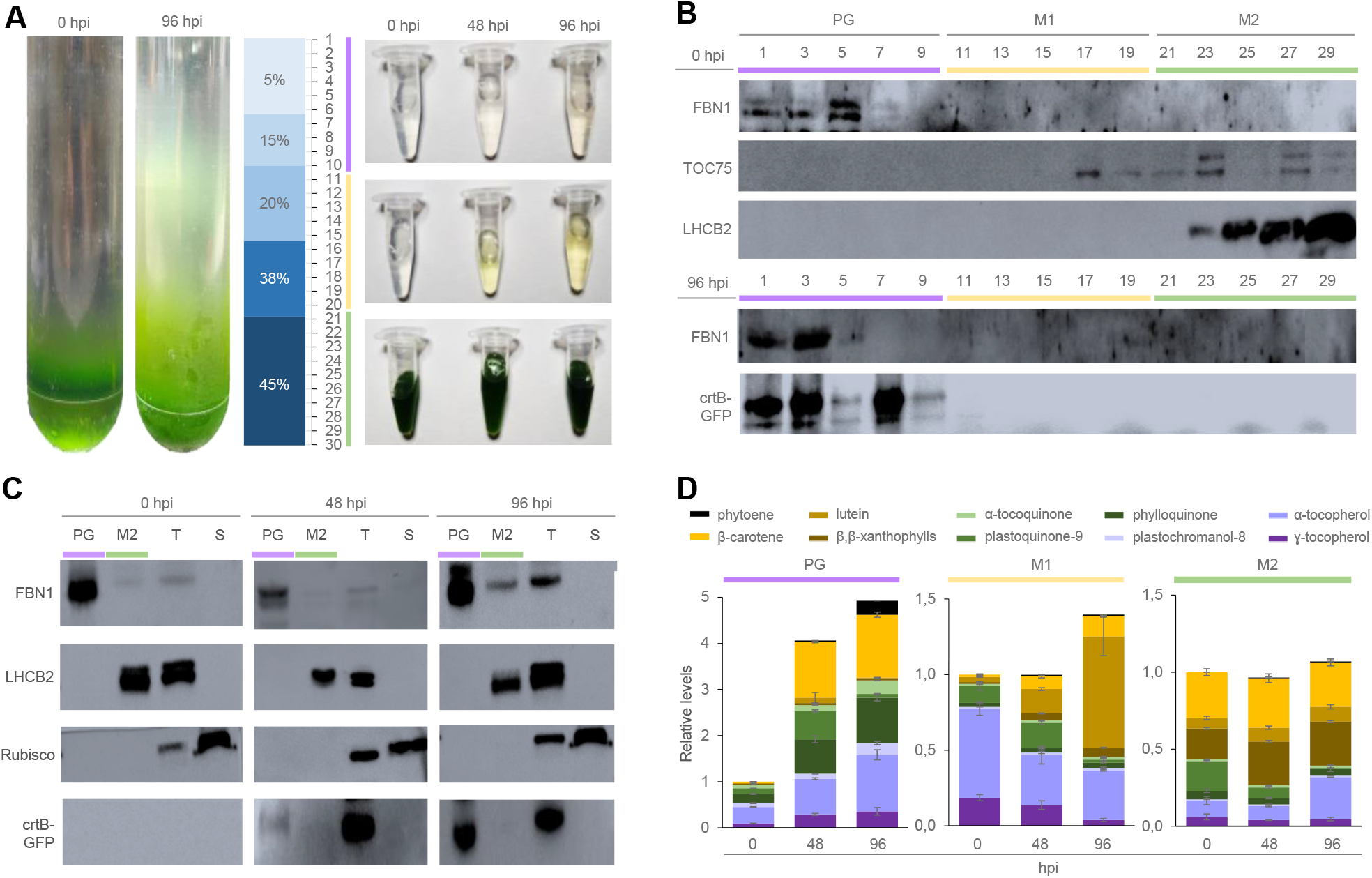
PG are a major site for the accumulation of isoprenoids in artificial leaf chromoplasts. Samples from *N. benthamiana* leaves agroinfiltrated with crtB-GFP constructs were collected at different timepoints. (A) Fractionation of plastidial membranes by flotation ultracentrifugation in a discontinuous sucrose gradient. The blue bar represents the position of layers with different sucrose concentrations (%) and the numbers in the right mark the fractions used for immunoblotting and HPLC analyses. Representative pictures of the samples after ultracentrifugation (left) and the collected fractions expectedly containing thylakoid membranes (M2, in green), envelope membranes (M2, in orange) or PG (in purple) are shown. (B) Immunoblot analysis of the fractions shown in (A) with antibodies against proteins from different plastid membranes. (C) Immunoblot analysis of the indicated fraction pools together with total plastid (T) or stromal (S) protein extracts (T). (D) Levels of the indicated isoprenoids in the indicated fraction pools. Data are represented relative to the levels in GFP samples and correspond to the mean and standard deviation of n=3 different leaves.

### Active crtB enzymes are located in the PG of artificial chloroplasts

PG are plastidial structures with a lipoprotein membrane monolayer that derive from thylakoids and are highly dynamic in size (Van Wijk and Kessler, 2017). They typically proliferate in response to environmental and developmental cues linked to a reorganization and disassembly of thylakoids such as excess light and senescence in leaves or ripening in fruits. In photosynthetic plastids (i.e. chloroplasts), PG harbor prenyl lipids such as chromanols and quinones and some of their biosynthetic enzymes such as VTE1 and NDC1 (Figure 1), whereas their main role in fruit and flower chromoplasts appears to be the biosynthesis, storage and degradation of carotenoids (Van Wijk and Kessler, 2017; Shanmugabalaji et al., 2022; Michel et al., 2021). Some plant PSY isoforms have been found to localize in PG using fluorescent protein tags in transient expression assays (Shumskaya et al., 2012). Using the same approach, we observed a full overlapping of fluorescent signals corresponding to crtB-GFP and the inducible PG marker FBN7a-RFP in tobacco chloroplasts (Figure 7), suggesting that the bacterial crtB might also be a PG-associated protein. In agreement with this conclusion, the crtB-GFP protein was found in the PG fractions obtained from the fractionation of plastidial membranes using flotation density centrifugation (Figure 6B-C). Furthermore, the observation that phytoene is mainly detected in these fractions (Figure 6D) strongly supports the conclusion that the crtB-GFP protein associates with PG in an enzymatically active form. This is in contrast to that observed in crtB-overexpressing tomato lines, in which crtB was associated to thylakoid-derived membranes of ripe fruit chromoplasts but it was not detected in the PG-rich fractions that contained most of the chromoplast phytoene content (Nogueira et al., 2013). Endogenous PSY1 (the tomato fruit isoform) was also absent from PG and mostly located in stromal fractions (Nogueira et al., 2013). Another difference between tobacco leaf (artificial) and tomato fruit (natural) chromoplasts was the localization of carotenoid products downstream of phytoene. Abundant carotenoids in tomato fruit such as lycopene and β-carotene were found at very low levels in PG fractions and mainly accumulated in the fractions corresponding to our M1 pool (Nogueira et al., 2013), while crtB-triggered leaf chromoplasts showed a preferential enrichment of β-carotene in PG and lutein in the M1 pool (Figure 6D). It is therefore likely that distribution of enzymes and metabolites changes between different chromoplast types, e.g., those developed in different plant tissues or/and species (Lado et al., 2015). Consistent with this conclusion, phytoene was most abundant in the PG-specific fraction of red pepper fruit chromoplasts, but also in a pepper-specific fibrillar fraction that also contained PG (Berry et al., 2019). The fibrillar fraction also contained most downstream carotenoids and α-tocopherol, as well as high levels of PSY that was less active than that found in membrane and stroma fractions (Berry et al., 2019). Interestingly, a different study found that carotenoid biosynthetic enzymes such as ζ-carotene desaturase (ZDS), lycopene β-cyclase (LCYB) and two β-carotene-hydroxylases (BCH) as well as the chromanol biosynthetic enzyme VTE1 (Figure 1) were present in the PG proteome of red pepper fruit chromoplasts (Ytterberg et al., 2006). Regardless of species and organ particularities, the results of these studies suggest a dynamic localization of carotenoid enzymes and products in different subplastidial membranes and subcompartments, with PG being one of major relevance in chromoplasts for the accumulation of these lipophilic metabolites.

**Figure 7.**
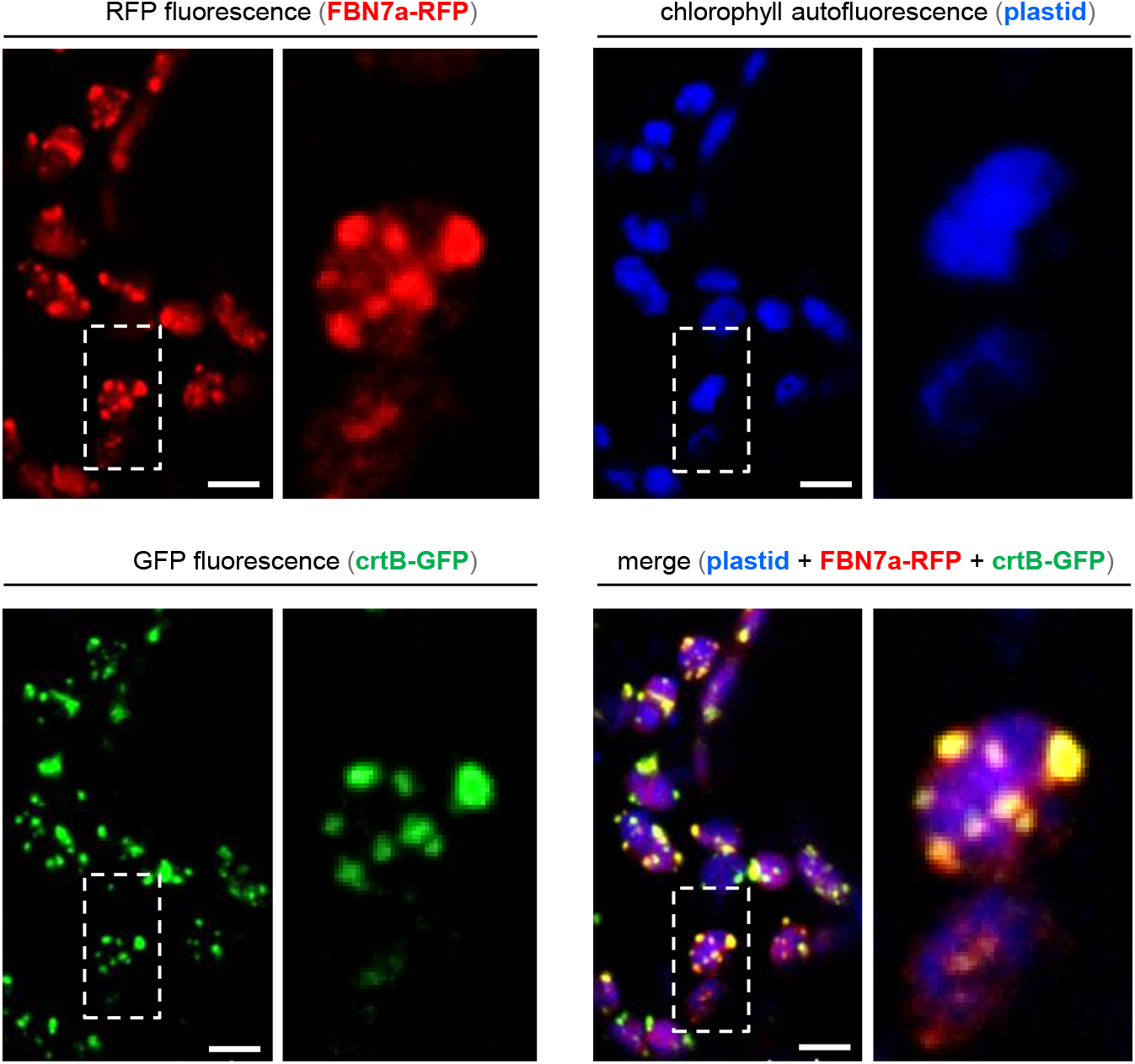
The crtB-GFP protein localizes to PG. *N. benthamiana* leaves were co-agroinfiltrated with constructs encoding the PG marker protein FBN7a-RFP and crtB-GFP and samples were collected at 96 hpi for confocal microscopy analysis. Fluorescence from FBN7a-RFP is shown in red, that from crtB-GFP in green, and chlorophyll autofluorescence in blue. Bar size, 5 µm. The dotted box marks the position of the image magnified on the right.

### High light pre-treatment results in improved levels of isoprenoid vitamins and a faster transition from chloroplasts to chromoplasts in crtB-inoculated leaves

Based on the data described above, we hypothesized that stimulating PG proliferation might further improve the accumulation of isoprenoid vitamins in crtB-induced leaf chromoplasts. To test this possibility, we used high light to promote PG development (Lundquist et al., 2013; Espinoza-Corral et al., 2021, 2019; Van Wijk and Kessler, 2017) and then induced artificial chromoplast differentiation with crtB (Figure 8). Whole *N. benthamiana* plants were acclimated for 3 days to white light (W) of either low or high intensity (50 and 500 μmol photons·m^−2^·s^−1^, respectively, herein referred to as W50 and W500) and then used for analysis of light curves to confirm the photoacclimation status of the plants (Figure 8A). As expected (Ralph and Gademann, 2005), W500-exposed plants showed lower photosynthetic rate (α) and maximum electron transport rate (ETRm) compared to those under W50 (Figure 8A). After the 3-day photoacclimation period, leaves were agroinfiltrated with crtB constructs and plants were incubated under normal growth conditions (W50) for 4 days. Estimation of chloroplast-to-chromoplast differentiation rate based on the decay of the photosynthetic parameter ɸPSII (Llorente et al., 2020) showed that pre-treatment with high light resulted in a faster chromoplastogenesis (Figure 8B). At 96 hpi, the overexpression of crtB resulted in substantially higher amounts of total carotenoids, tocopherols and phylloquinone in leaves pre-exposed to W500 (Figure 8C), hence validating our hypothesis. Exposure to high light may lead to the production in chloroplasts of reactive oxygen species (ROS) such as singlet oxygen (^1^O_2_), superoxide anion (O_2_^-^), hydroxyl radical (OH^-^) and hydrogen peroxide (H_2_O_2_) if photons that are not used for photosynthesis are not safely dissipated (Muñoz and Munné-Bosch, 2018; Li and Kim, 2022) ROS can oxidize nearby macromolecules and damage the structural and functional integrity of the chloroplast, but they are also involved in so-called retrograde signaling, i.e., communication of chloroplasts with the nucleus to readjust gene expression and restore redox homeostasis. To test whether the acceleration of chromoplast differentiation in W500-exposed tissues was associated to a higher production of ROS, we stained leaves with 3,3’-diaminobenzidine (DAB, which produces a brown precipitate when oxidized by H_2_O_2_) and nitrotetrazolium blue chloride (NBT, which reacts with O_2_^-^ to form a dark blue insoluble compound) (Figure 9). Under normal conditions, i.e., in plants grown under W50 before and after agroinfiltration, crtB-inoculated sections showed a strong up-regulation of ROS accumulation after 24 hpi (Figure 9), paralleling the onset of phytoene increase and ɸPSII decrease (Figure 3A). By contrast, GFP-agroinfiltrated sections of the same leaves showed a much lower ROS increase (Figure 9), suggesting that the enhanced ROS boost in crtB areas is not merely a consequence of agroinfiltration but it is most likely linked to the chromoplastogenesis process. As expected, leaves pre-exposed to W500 showed higher levels of ROS than control leaves before agroinfiltration, at 0 hpi (Figure 9). This was particularly evident in the case of O_2_^-^ (NBT staining). Then, levels dropped in GFP-agroinfiltrated sections to reach the basal levels detected in leaves not pre-acclimated to W500 whereas it further increased in crtB-agroinfiltrated areas (Figure 9). Based on these data, we concluded that the higher ROS levels present in W500-acclimated leaves might be associated to the acceleration of chloroplast-to-chromoplast differentiation that takes place after agroinfiltration with crtB. It is remarkable that the presence of high ROS levels in crtB sections occurs despite the progressive increase in powerful antioxidants such as tocopherols and carotenoids (Figure 6). However, this is in agreement with the observation that ROS accumulation and oxidative stress normally intensifies when chromoplasts naturally differentiate from chloroplasts during fruit ripening, possibly due to a gradual loss in the capacity of antioxidant enzymes to scavenge the excessive free radical production (Muñoz and Munné-Bosch, 2018; González-Gordo et al., 2019).

**Figure 8.**
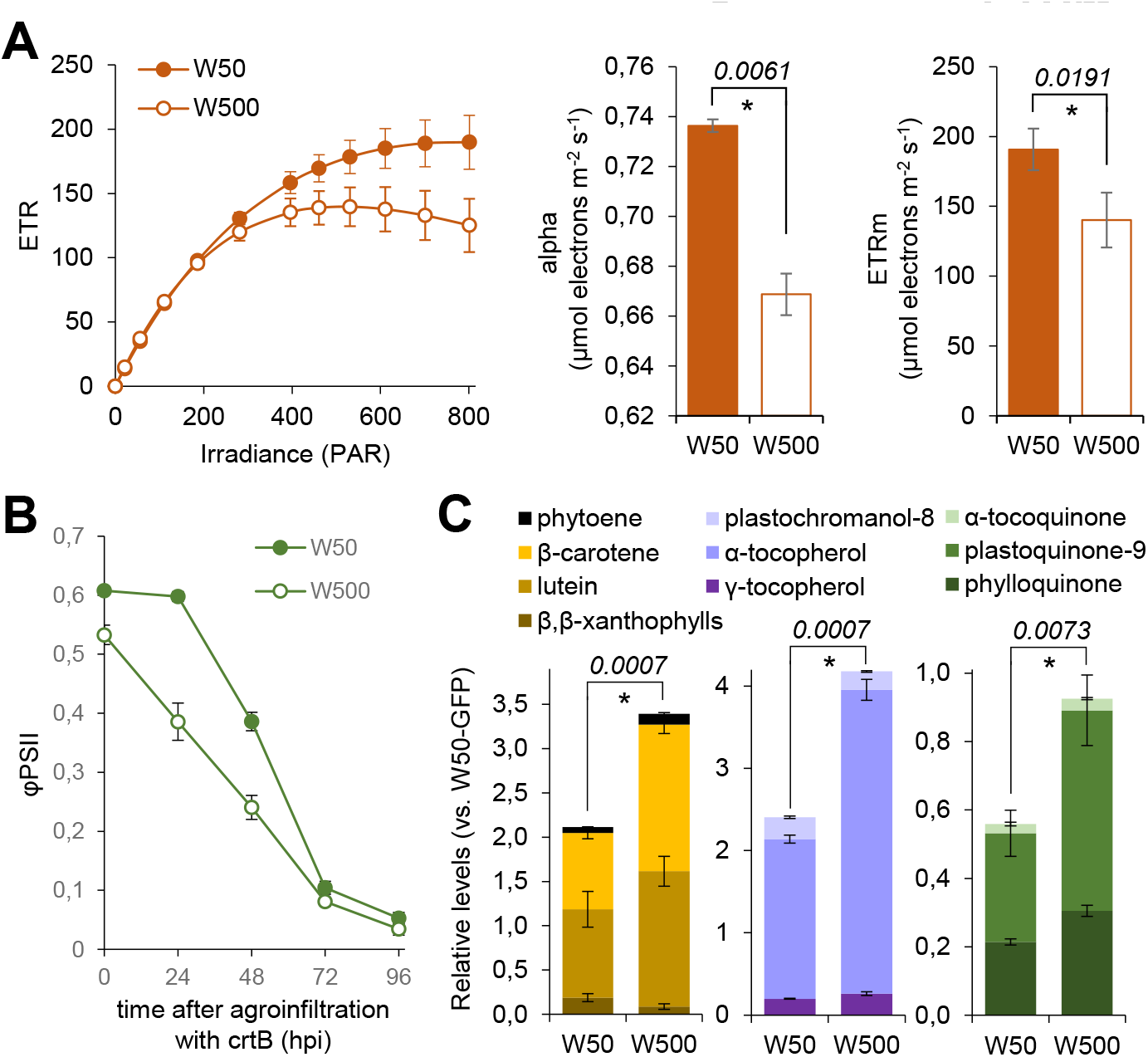
High light promotes chromoplastogenesis. *N. benthamiana* plants grown under normal white light conditions (W50) were exposed for 3 days to 10-fold higher light intensity (W500) or kept under W500. Then, leaves from the two sets of plants were agroinfiltrated with crtB constructs. (A) Photosynthetic electron transport rates (ETR) vs irradiance curve (left) and derived parameters: alpha (photosynthetic rate in the light-limited region of the curve) and ETRm (maximum ETR) of leaves from plants exposed to either W50 or W500 before agroinfiltration (0 hpi). (B) Evolution of φPSII values in crtB-agroinfiltrated leaves from plants pre-exposed to either W50 or W500. (C) Levels of the indicated isoprenoids at 96 hpi. In all plots, data correspond to the mean and standard deviation of n=3 different samples. Numbers in italics indicate *p* values of Student *t*-test analyses and asterisk marks statistically significant differences (*p* ≤ 0.01).

**Figure 9.**
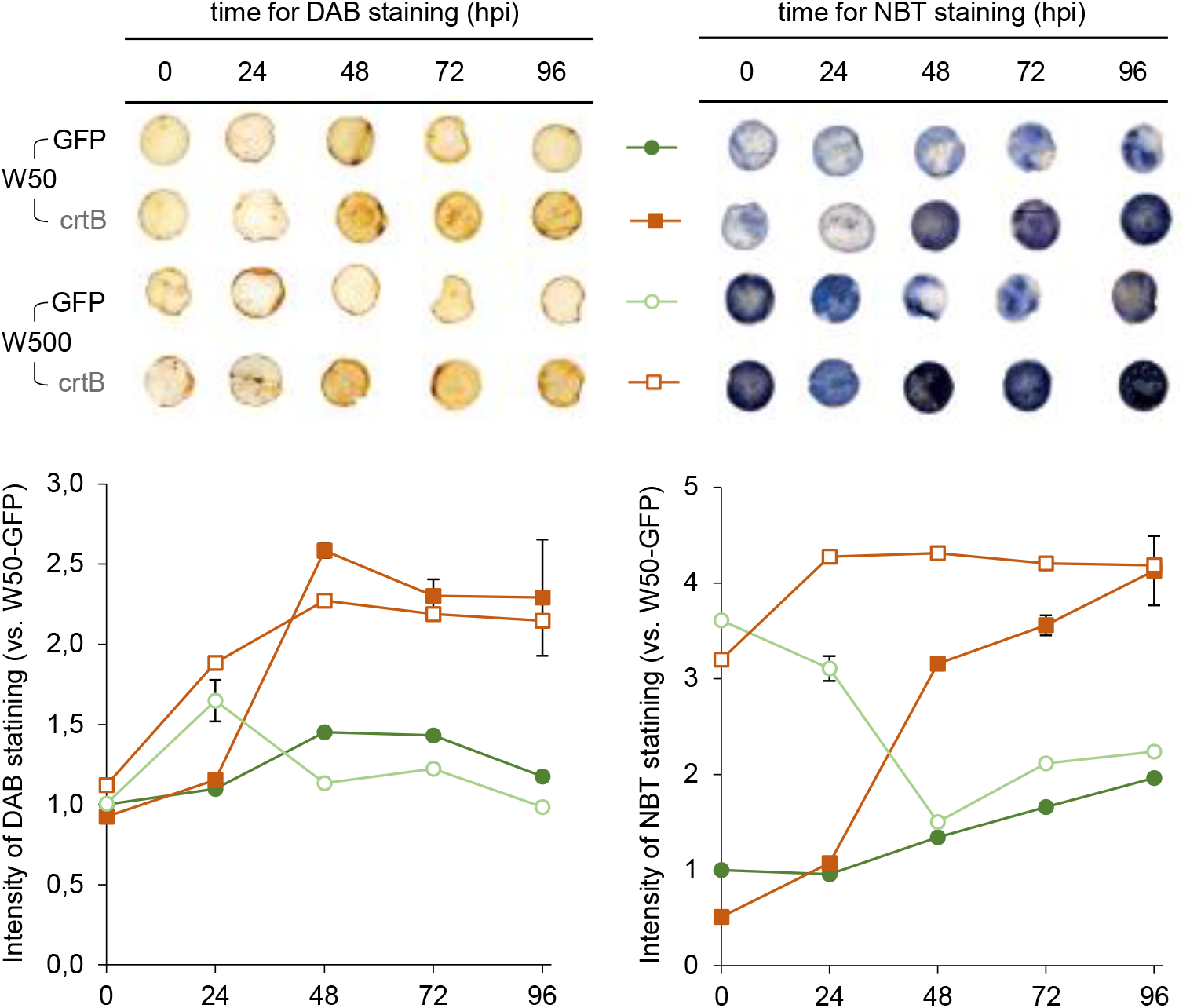
ROS production is associated to chromoplastogenesis. *N. benthamiana* plants grown under normal white light conditions (W50) were exposed for 3 days to 10-fold higher light intensity (W500) or kept under W500. Then, leaves from the two sets of plants were agroinfiltrated with either GFP or crtB constructs and samples were collected and used for staining with DAB or NBT at the indicated timepoints. Representative images of stained leaf samples (circles) are shown. Plots show quantitative data (mean and standard deviation) from n=3 independent samples.

### crtB-triggered chromoplastogenesis as a model system to understand chloroplast to chromoplast differentiation

The differentiation of chloroplasts into chromoplasts is a very important process for plants (as it provides flower and fruit colors to attract animals for pollination and seed dispersal) but it also has an applied interest (as it typically results in nutritional enrichment of the tissue in health promoting carotenoids and other isoprenoids). Yet our scarce knowledge of how chromoplastogenesis occurs has prevented to efficiently manipulate this process for food biofortification, among other applications (Torres-Montilla and Rodriguez-Concepcion, 2021). In part, our fragmented understanding of how chloroplasts become chromoplasts is due to the difficulties associated with the study of natural chromoplastogenesis, as this process is normally linked to developmental processes such as petal or root development and fruit ripening. By uncoupling chromoplast differentiation from other developmental processes, our crtB-dependent system producing artificial chromoplasts from leaf chloroplasts is providing new insights. Our data indicate that protein import and PG development are important for chromoplast differentiation, but their specific roles may change in different plant systems. For example, SP1 and TOC75 appear to be key to regulate plastid protein import for chromoplastogenesis in tomato fruit (Ling et al., 2021) but not in our artificial leaf system, which instead might rely on TOC33 (Figure 3). In the case of PGs, we conclusively show that they are not mere reservoirs of carotenoids but active production sites (Figure 6), even though the specific carotenoid enzymes and products they host changes in other systems and even among different plant species that naturally differentiate PG-containing chromoplasts during fruit ripening (Ytterberg et al., 2006; Nogueira et al., 2013; Berry et al., 2019). Most importantly, our synthetic system has also unveiled that some of the features originally believed to be consequences of chromoplast differentiation are actually requirements for chloroplasts to become chromoplasts. Thus, up-regulated carotenoid production together with null or reduced photosynthetic activity are necessary to trigger the chromoplastogenesis process (Llorente et al., 2020). In the work reported here we demonstrate that the production of ROS is also a trigger of the differentiation process (Figure 9). While it remains unclear whether ROS production is a cause or/and a consequence of chromoplastogenesis during the fruit ripening process, it has been reported that oxidative stress stimulates the conversion of fruit chloroplasts into chromoplasts, which in turn facilitates the synthesis and accumulation of antioxidants such as carotenoids (Fanciullino et al., 2014; Corpas et al., 2018). Correlative evidence for a possible role of redox signaling in promoting fruit ripening is also available ((Fanciullino et al., 2014; Muñoz and Munné-Bosch, 2018; Corpas et al., 2018; Decros et al., 2019; Li and Kim, 2022). Redox sensitive systems rather than ROS appear to regulate both the expression of genes of the carotenoid pathway and the activity of redox-dependent enzymes (Fanciullino et al., 2014). We therefore conclude that, similar to that already described for the down-regulation of photosynthesis and the accumulation of carotenoids, the production of ROS and likely the subsequent redox changes are an integral part of the chromoplast differentiation process and not just a consequence.

The generation of chromoplasts is one of the most promising strategies for the biofortification of green leaves despite the low number of tools available to control the process. In this work, we have shown that crtB is a powerful tool to transform leaf chloroplasts into artificial chromoplasts in a process that involves a reduction in photosynthetic activity, proliferation of PG enriched in isoprenoid vitamins such as β-carotene (pro-vitamin A), α-tocopherol (vitamin A) and phylloquinone (vitamin K1), and enhanced production of ROS as chloroplasts lose their functionality. Despite this being a synthetic system, it shares general features with natural chromoplastogenesis processes and hence it represents a unique model to better understand how a chromoplast is made. This knowledge should contribute to creating artificial sinks to improve both the production and the storage of carotenoids and other isoprenoid phytonutrients in green tissues but also all other sorts of plant-derived foods.

## Materials and methods

### Plant material, growth conditions and photosynthetic measurements

*Nicotiana benthamiana* and *Arabidopsis thaliana* plants were grown as described previously (Llorente et al., 2016; Majer et al., 2017; Andersen et al., 2021). For the intense light experiments, plants were grown under standard conditions and then moved to an ARALAB 600 growth chamber where they were maintained for three days in two different light conditions: 50 μmol photons·m^−2^·s^−1^ (named W50) and 500 μmol photons·m^−2^·s^−1^ (W500) under a long day (16h light and 8h dark) photoperiod. After agroinfiltration, plants were all maintained in the W50 condition. Photosynthetic measurements were performed as described (Llorente et al., 2020). Light curves (ETR vs Irradiance) and derived parameters (alpha and ETRm) were generated as described (Morelli et al., 2021) with 11 incremental steps of actinic irradiance (E; 0, 21, 56, 111, 186, 281, 396, 461, 531, 611, 701, 801 μmol photons·m^−2^·s^−1^).

### Gene constructs

The crtB versions used for this study were obtained as described (Llorente et al., 2020). To generate inducible PG markers, cDNA obtained from *A. thaliana* seeds and leaves was used to amplify sequences encoding full-length VTE1 and FBN7a, respectively. PCR products were cloned using the Gateway system first into a pDONR-207 plasmid and then into plasmid pGWB504 provided with an RFP fluorescent tag (Nakagawa et al., 2007). The generated constructs (p35S:VTE1-RFP and p35S:FBN7a-RFP) were then used as templates for PCR-based amplification of the RFP-fused proteins. The two amplicons were then cloned into pDONR207 and then into the β-estradiol-inducible vector pER8 to obtain pXVE:VTE1-RFP and pXVE:FBN7a-RFP. Similarly, cDNA sequences encoding full-length TIC40 and TOC64 envelope proteins were amplified and cloned first into a pDONR-207 plasmid and later into the β-estradiol-inducible vector pMDC7 (provided with a CFP fluorescent tag) to obtain pXVE:TIC40-CFP and pXVE:TOC64-CFP constructs. Primers for these clonings are listed in Table S3. Viral vectors TuMV and TuMV-crtB (Llorente et al., 2020) were kindly provided by José Antonio-Daròs.

### Transient expression assays

Agroinfiltration assays were carried out as described (Andersen et al., 2021) using cells from fresh *Agrobacterium tumefaciens* GV3101 cultures transformed with the appropriate constructs at an optical density at 600 nm of 0.5 in infiltration buffer (10 mM MES, 10 mM MgCl_2_, 150 µM acetosyringone, pH=5.5). Cultures were mixed in identical proportions when agroinfiltrating several constructs. For the induction of fluorescent proteins, a solution of 17-β-estradiol (Sigma) was prepared by diluting the compound to a concentration of 100 µM in water and 0.05% Tween-20 and applied with a fine brush on the leaf 6 hours after the infiltration. Inoculation of viral vectors was carried out as described (Llorente et al., 2020).

### Chloroplast isolation and membrane fractionation

Whole chloroplasts were isolated from leaves blended into a Waring mixer with 200 ml of HB buffer (20 mM Tricine/KOH pH=8.4, 450 mM sorbitol, 10mM EDTA, 10 mM NaHCO_3_, 1 mM MgCl_2_, 5 mM sodium-ascorbate, 1 mM PMSF). The mixture was then filtered through 2 layers of cheesecloth and one layer of miracloth at 4°C and then centrifuged for 10 min at 750 x*g*. The pellet was resuspended in 2 ml of HB buffer to measure chlorophyll concentration as described (Shanmugabalaji et al., 2013). The volume was then increased to 50 mL with HB buffer and the samples were centrifuged again at 4°C at 1250 x*g* for 2 min. The pellet was resuspended in 0.6 M sucrose solution in TED buffer (500mM Tricine/KOH pH=7.5, 20mM EDTA, 20mM DTT) to reach a final chlorophyll concentration of 2 mg/ml. Chloroplast suspension was then diluted with 3 volumes of TED buffer, manually homogenized, and then centrifuged for 1 h at 100.000 x*g.* The upper phase (stroma) was removed and frozen for further analysis, and the pellet was resuspended into 45% sucrose in TED buffer at a concentration of 3 mg/ml of chlorophyll. The suspension was homogenized again in a glass potter and the resulting resuspended membranes were pipetted into a polycarbonate UltraClear SW28 tube. A discontinuous gradient of sucrose in TED buffer was then layered on top of the membranes (6 ml 38% sucrose, 6 ml 20% sucrose, 4 ml 15% sucrose, 6 ml 5% sucrose). The obtained gradient was then centrifuged at 100.000 x*g* for 16 h. Single fractions were then collected and stored for further analysis.

### Protein extraction and immunoblot analyses

Total protein extraction from freeze-dried leaf material was carried out as described (Shanmugabalaji et al., 2018). For the isolation of proteins from chloroplast fractions, 200 µl samples were collected and precipitated by adding 4 volumes of acetone and incubating the samples at -20°C for 30 min. SDS-PAGE and immunoblot analyses were performed as described (Shanmugabalaji et al., 2018). After protein transfer, the nitrocellulose membranes were stained with amido black 10B staining (Sigma) to detect the Rubisco protein as an indicator of successful transfer and equal loading. Primary antibodies for Arabidopsis TOC33, TOC75, TOC159, FBN1a, FBN2, FBN4 and VTE1 were previously available in the lab (Vidi et al., 2006; Rahim et al., 2009; Shanmugabalaji et al., 2013). Commercial antibodies were also used against GFP (Sigma) and plant Rubisco, LHCB2, PsaD, PsbA, PetC and TIC40 proteins (Agrisera). HRP-conjugated anti-rabbit (Millipore) or anti-mouse (Sigma) IgG antibodies were used as secondary antibodies and chemiluminescent signals were detected using ECL Plus Western Blotting Detection Reagents (Pierce) and a GE Amersham Imager 600. Band intensities were quantified using ImageJ software (Schindelin et al., 2012).

### Metabolite analyses

Leaf carotenoids, chlorophylls and tocopherols were extracted and analyzed by HPLC-DAD-FLD as described (Barja et al., 2021). The rest of isoprenoid metabolites were separated and quantified by UHPLC-QTOF-MS as described (Spicher et al., 2016; Martinis et al., 2011). Phytoene levels were measured using both methods. In the case of the UHPLC system, an Acquity UHPLC coupled to a Synapt G2 (Waters) was used as follows: the separation was performed on an Acquity BEH C18 (100x2.1 mm, 1.7 µm particle size) maintained at 70°C. Mobile phases were H_2_O + 0.1% NH_4_OH (A) and methanol + 0.1% NH4OH (B). The flow rate was 0.4 ml/min and a program consisting of a gradient from 80% B to 100% B in 6 min followed by a hold at 100% for 3 min was applied. Under these conditions, phytoene eluted at 8.37 min. Detection was performed at 285 nm with a diode array detector and by electrospray positive ionization using a scan range of 50-600 Da (with enhanced duty cycle set to m/z 545.51). The system was controlled by Masslynx 4.1. Absolute quantification was achieved by external calibration using a phytoene standard (CaroteNature).

### Microscopy

Subcellular localization of fluorescent proteins was performed by direct examination of agroinfiltrated leaf tissue at the indicated time points with a Leica TCS SP8-MP Confocal Laser Scanning Microscope. GFP fluorescence was detected using a BP515-525 filter after excitation at 488 nm while CFP was detected after excitation at 440 nm. Chlorophyll autofluorescence was detected using a 610-700 nm filter after excitation at 568 nm while the RFP signal was detected after excitation at 532 nm laser line and detected at 588 nm. Transmission electron microscopy was carried out as described (Llorente et al., 2020). Subcellular structures were quantified by using the ImageJ software (Schindelin et al., 2012).

### ROS histochemical staining

*N. benthamiana* leaf disks were placed inside a syringe body to be infiltrated with a solution of 1 mg/ml 3,3’-diaminobenzidine (DAB, Sigma) by moving the plunger up and down 10-15 times. Similarly, leaf discs were infiltrated with a 0.1% solution of nitrotetrazolium blue chloride (NBT, Sigma) in 50mM phosphate buffer (pH=7.8) and incubated at room temperature in the dark for 2 h. After incubation with DAB or NBT, leaf disks were bleached by incubating in a 3:1:1 solution of ethanol:lactic acid:glycerol for 8 h and then mounted in 60% glycerol on transparent paper and scanned for further analysis. Intensity of DAB or NBT staining was quantified by using the ImageJ software (Schindelin et al., 2012).

### Transcript and phylogenetic analyses

Transcript abundance was deduced from previously published RNAseq data (Llorente et al., 2020). Homologue genes were identified with BLAST using *Arabidopsis* (Araport11) genes as query against tomato (ITAG4.0), pepper (*Capsicum_annuum*.ASM51225v2) and *N. benthamiana* (Niben261) genomes. Only sequences with a sequence identity of 35% or higher were selected and concatenated in a single fasta file to be aligned with Guidance (Sela et al., 2015) using ClustalW. To create a phylogenetic tree, iqtree (Nguyen et al., 2015) was run to select the best model fitting the aligment. Once the model was selected, iqtree was run for a second time with that model. Genes were annotated one by one based on their clustering with Arabidopsis. Those from tomato were validated in SolGenomics and used to annotate *N. benthamiana* genes. Normal clustering was 1:1:2 (tomato:pepper:*N. benthamiana*, respectively). All heatmaps were created in RStudio using pheatmap package (Kolde, 2019).

## Acknowledgments

We thank José Antonio Daròs for providing viral vectors. This work was funded by grants from Spanish MCIN/AEI/10.13039/501100011033 and European NextGeneration EU/PRTR and PRIMA programs to MR-C (PID2020-115810GB-I00 and UToPIQ-PCI2021-121941). MR-C is also supported by CSIC (202040E299) and Generalitat Valenciana (PROMETEU/2021/056). Work at University of Neuchatel was supported by the Swiss National Science Foundation (SNSF) grant 31003A_176191. LM and ST-M received predoctoral fellowships from La Caixa Foundation (INPhINIT program LCF/BQ/IN18/11660004) and MEC (FPU program, FPU16/04054), respectively. LM was also funded by an EMBO short-term fellowship (ASTF 8907) to do part of the work at the University of Neuchatel (Switzerland).

## Author Contributions

LM and MR-C designed the research; LM, ST-M, GG and VS performed research; LM, ST-M, GG and VS contributed new analytic and computational tools; LM, ST-M, GG, VS, FK and MR-C analyzed data; LM and MR-C wrote the paper.

---

**Figure S1.**
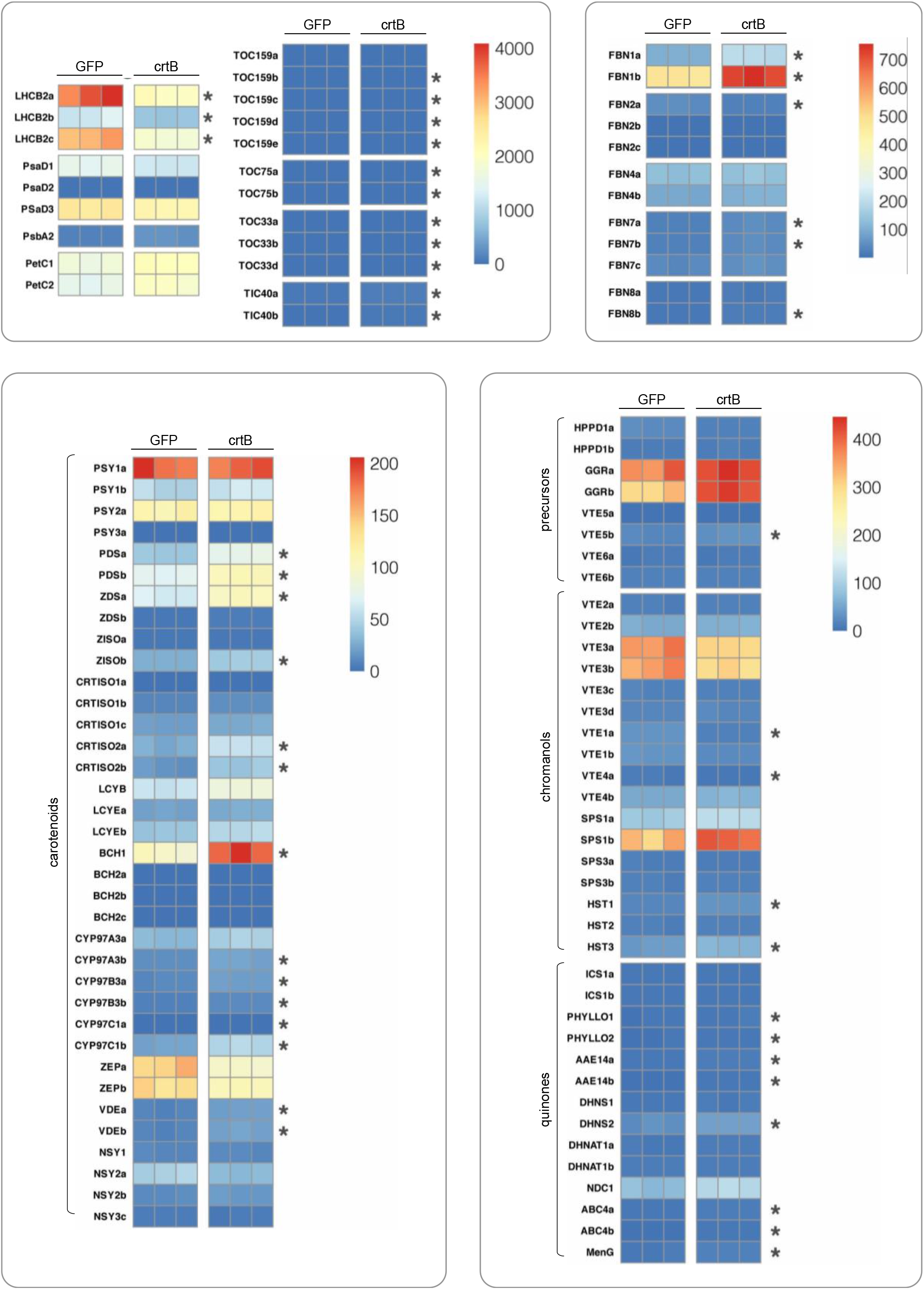
Transcript abundance of the indicated genes in GFP and crtB samples at 96 hpi. Data were retrieved from previously reported RNA-seq analysis (Llorente et al., 2020). Columns correspond to replicates. Transcript levels are shown as absolute values. Asterisks mark differentially expressed genes when comparing mean crtB vs. GFP values (DESeq2, FDR ≤ 0.05). Gene accessions are available in Tables S1 and S2.

**Table S1.**
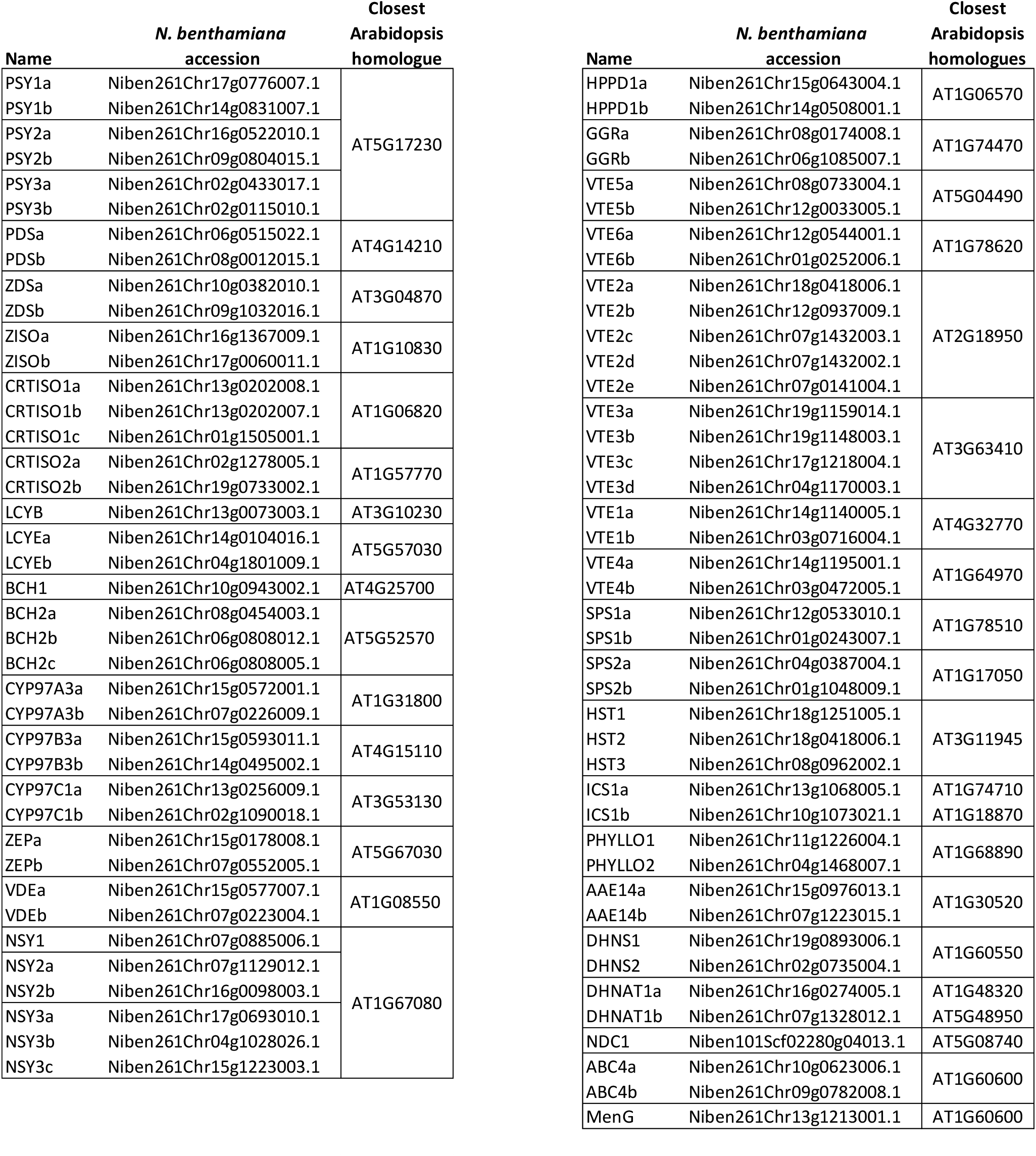
Accessions of genes encoding isoprenoid biosynthetic enzymes.

**Table S2.** Accessions of genes encoding genes for plastid proteins.

| Name | *N. benthamiana*<br>accession | Closest<br>Arabidopsis<br>homologues |
| --- | --- | --- |
| LHCB2a | Niben261Chr04g1775002.1 | AT2G05100 |
| LHCB2b | Niben261Chr19g0203003.1 |  |
| LHCB2c | Niben261Chr11g0348018.1 |  |
| PsaD1 | Niben261Chr13g1342002.1 | AT1G03130<br>AT4G02770 |
| PsaD2 | Niben261Chr05g0971027.1 |  |
| PsaD3 | Niben261Scf14940g0000002.1 |  |
| PsbA2 | Niben261Chr03g1621009.1 | ATCG00020 |
| PetC1 | Niben261Chr14g0161005.1 | AT1G78620 |
| PetC2 | Niben261Chr08g1132007.1 |  |
| TOC159a | Niben261Chr17g0812006.1 | AT4G02510 |
| TOC159b | Niben261Chr13g0349002.1 |  |
| TOC159c | Niben261Chr08g0076035.1 |  |
| TOC159d | Niben261Chr10g0004003.1 |  |
| TOC159e | Niben261Chr09g0431013.1 |  |
| TOC75a | Niben261Chr13g1023003.1 | AT3G46740 |
| TOC75b | Niben261Chr10g1172001.1 |  |
| TOC33a | Niben261Chr08g0566009.1 | AT1G02280 |
| TOC33b | Niben261Chr06g0969009.1 |  |
| TOC33d | Niben261Chr01g0946016.1 |  |
| TIC40a | Niben261Chr10g0714004.1 | AT5G16620 |
| TIC40b | Niben261Chr03g1913001.1 |  |
| FBN1a | Niben261Chr16g0529004.1 | AT4G04020 |
| FBN1b | Niben261Chr07g1400002.1 | AT4G22241 |
| FBN2a | Niben261Chr18g0995001.1 | AT2G35490 |
| FBN2b | Niben261Chr05g1310005.1 |  |
| FBN2c | Niben261Chr02g1180003.1 |  |
| FBN4a | Niben261Chr09g0283001.1 | AT3G23400 |
| FBN4b | Niben261Chr04g1138001.1 |  |
| FBN7a | Niben261Chr18g1220007.1 | AT2G42130<br>AT3G58010 |
| FBN7b | Niben261Chr13g0130005.1 |  |
| FBN7c | Niben261Chr08g0918006.1 |  |
| FBN8a | Niben261Chr14g1141015.1 | AT2G46910 |
| FBN8b | Niben261Chr03g0716005.1 |  |

**Table S3.**
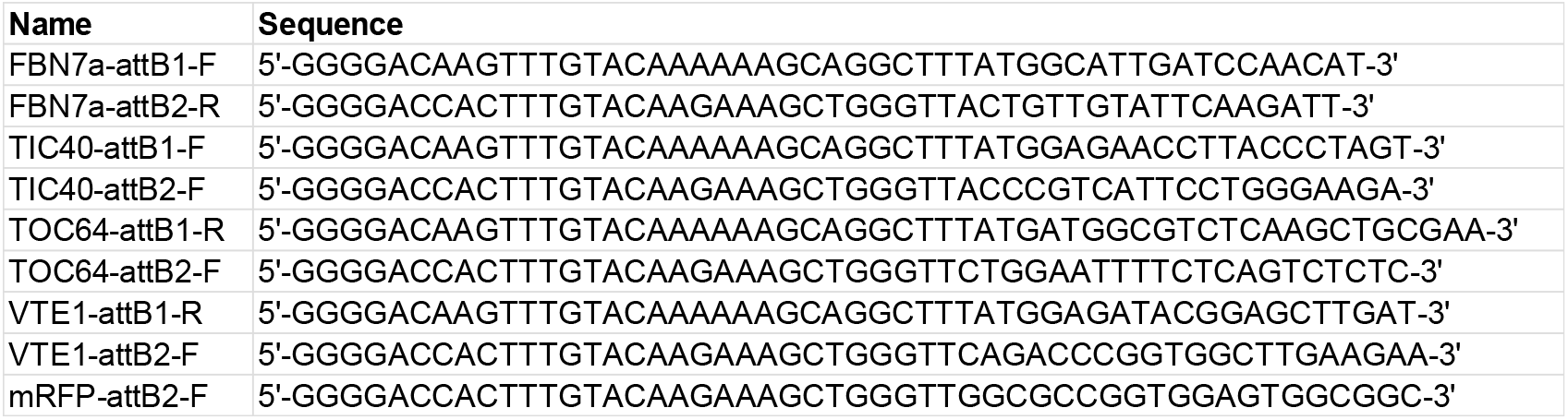
. Primers used in this work.

## Notes

### Competing Interest Statement

The authors have declared no competing interest.

